# Use of a visible reporter marker- *myb*-related gene in crop plants to minimize herbicide usage against weeds

**DOI:** 10.1101/455162

**Authors:** Radi Aly, Ran Lati, Vinay K. Bari, Jackline Abu-Nassar, Hanan Eizenberg

## Abstract

Weeds, a main threat to agricultural productivity worldwide, are mostly controlled by herbicides. To minimize herbicide usage by targeting it to weedy areas, we developed a new image-based methodology for robust weed detection that relies on manipulating the crop plant’s leaf hue, without affecting crop fitness. We generated transgenic tobacco (*Nicotiana tabacum* Xanthi) lines overexpressing anthocyanin pigment as a traceable marker that differentiates transgenes from the surrounding weeds at an early stage. Transformation with the anthocyanin *VlmybA1-2* gene produced purple-colored leaves. Subsequent gene silencing with vector p*TRV2*:*VlmybA1-2* significantly reduced anthocyanin pigments in tobacco leaves 40 days after agroinfiltration, with a concomitant reduction in *VlmybA1-2* transcript levels. Purple hue faded gradually, and there were no fitness costs in terms of plant height or leaf number in the silenced vs. non-silenced tobacco transgenes. These results could lead to a new sustainable weed-control method that will alleviate weed-related ecological, agricultural and economic issues.

## Introduction

Herbicides are the basis of most weed management, but the negative aspects associated with their application have put great pressure on farmers and regulators to reduce or ban their use (1). By targeting herbicide application exclusively to weedy areas, precise weed management (PWM) methods are a promising means of minimizing herbicide usage without affecting crop yield or quality (2). To implement PWM methodologies, it is essential to identify and distinguish between weeds and crops, considered to be the bottleneck in the development of any PWM method (3).

RGB cameras are the prime tool for weed detection and classification due to their low cost and high viability (4). Since crops and weeds share a similar “green” hue, the RGB-based methods utilize advanced shape, texture and size algorithms for classification (5). Nonetheless, adequate results have mainly been achieved under specific controlled conditions and early crop stages, whereas under non-controlled field conditions, they have not been as robust (6). An alternative classification approach was suggested by Lati et al. (7) that used genetically modified tomato plants with overexpression of anthocyanin (germplasm AN-113) which expressed purple leaves throughout their growth. The sharp hue difference between the purple-leaved tomato and the surrounding “green” weeds yielded absolute declassification rates under various growth stages and illuminations. However, in our previous approach Lati et al. (7), we observed potential negative impact of anthocyanin overexpression in vegetative and fruit tissues of tomato plants. Since weed control is most important at the crop’s early growth stages, before its canopy is well developed and the plants become strong competitors, we assumed that overexpression of anthocyanin can be limited to this early stage to avoid potential reduction in crop fitness.

*MybA* genes are known to be involved in the regulation of anthocyanin biosynthesis in several species (8–10). To limit the crop’s overexpression of anthocyanin to early growth stages, the virus-induced gene-silencing (VIGS) system (11), which has become a powerful tool for the analysis of gene function, reverse genetics (12) and gene silencing, can be used (13–15). Here, we overexpressed anthocyanin as a visible marker in the crop plant at an early growth stage for robust weed classification, then decreased anthocyanin biosynthesis during crop development using the VIGS system. We propose this method as an ideal solution for obtaining high weed-detection rates at early growth stages while avoiding potential reductions in crop performance.

### Results and Discussion

To obtain transgenic crops overexpressing anthocyanin at early growth stages, we integrated a method suggested by Orzaez et al. (12), a visual reporter system for VIGS in tomato fruit based on anthocyanin accumulation with another method suggested by Tian et al. (16) that used a modified Tobacco rattle virus (TRV)-VIGS system to evaluate the degree of silencing of a target gene. Here, we overexpressed *VlmybA1-2* encoding anthocyanin pigment in transgenic tobacco lines (figure 1, I): (B and C), resulting in purple-leaf phenotypic features. To make these transgenic lines useful for precise weed control while avoiding fitness costs, we investigated whether we can reduce the levels of anthocyanin at later growth stages using the p*TRV2*:*VlmybA1-2* vector as a VIGS system. A mixture of *Agrobacterium* cultures containing (*pTRV1*) and p*TRV2*:*VlmybA1-2* were agroinfiltrated 1 week after transplantation of transgenic tobacco plants expressing anthocyanin to pots. About 40 days after agroinfiltration, gene silencing was successful: anthocyanin pigments were significantly reduced in tobacco leaves treated with p*TRV2*:*VlmybA1-2* as compared to the non-silenced control line with an empty vector (pTRV1:pTRV2) (Figure 1, I). To study transcriptional changes in the anthocyanin gene, RT-PCR analysis was performed: transcript levels of *VlmybA1-2* were substantially reduced in p*TRV2*:*VlmybA1-2*-agroinfiltrated leaves relative to tobacco leaves infiltrated with empty vector (Figure 1, II). Accordingly, the silenced p*TRV2*:*VlmybA1-2* lines showed completely “purple”-colored leaves in the first 5 weeks after agroinfiltration (Figure 2). Then, the purple hue gradually faded; by the end of the study, there was no significant difference between the green areas of the silenced line and the WT. In contrast, the tobacco lines agroinflitrated with empty vector, showed strong purple color throughout the entire study (Figure 2). In addition, Figures 3a and 3b show that during the study, the silenced lines showed slight fitness cost in terms of plant height or leaf number. In contrast, the control lines with the empty vector were completely stunted and showed a significant reduction in leaf number and height compared to the WT plants and the silenced lines (Figure 3).

**Figure 1.**
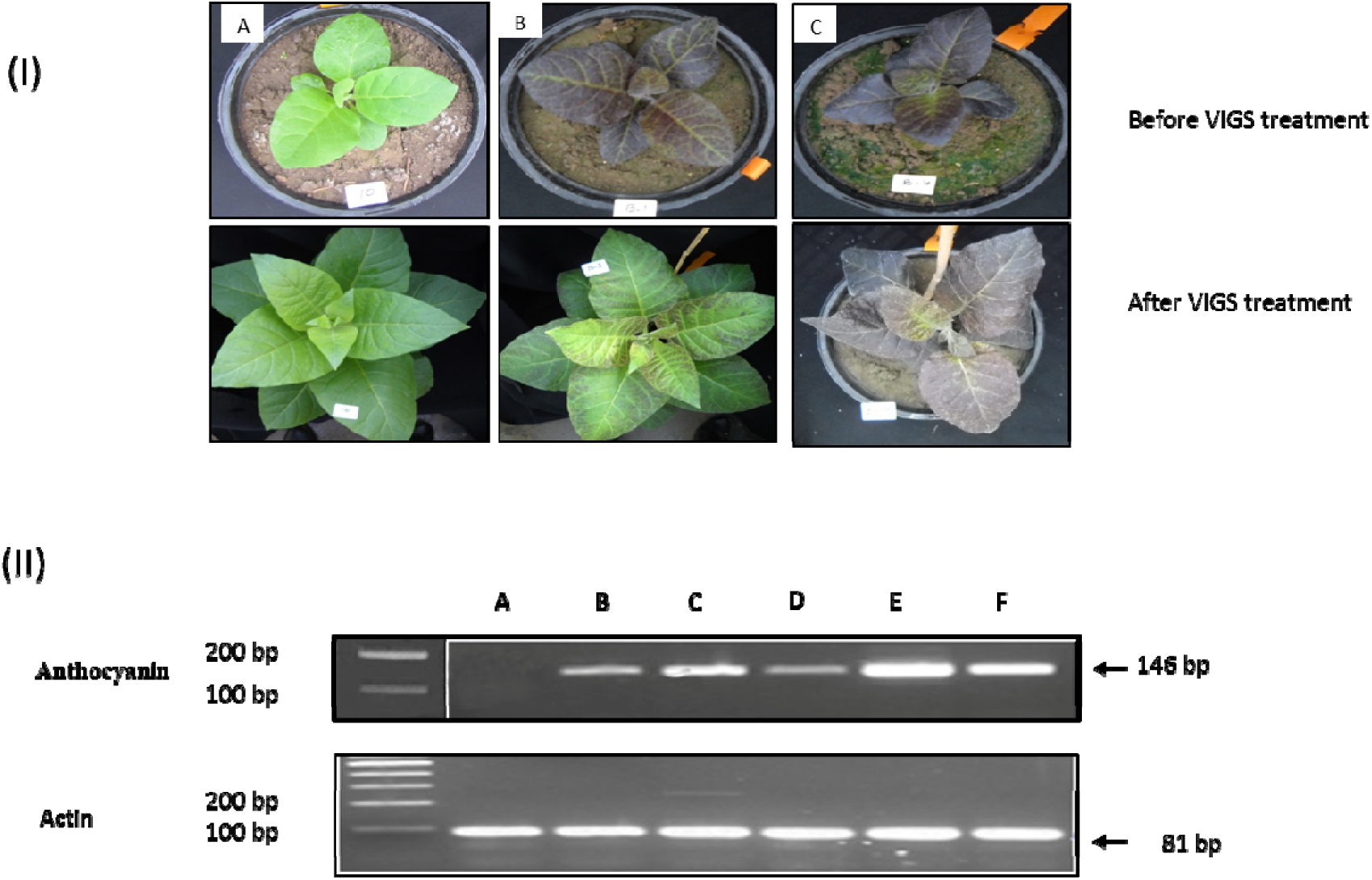
Overexpression, suppression and abundance of anthocyanin gene *VlmybA1-2* in *N. tabacum* (Xanthi) plants in two representative lines (B and C). (**I**) Upper row: WT (A) and transgenic *N. tabacum* (Xanthi) seedlings overexpressing anthocyanin pigments before VIGS treatment (B and C). Lower row: suppression of anthocyanin mRNA in *N. tabacum* (Xanthi) leaves via *pTRV2*:*VlmybA1-2* (B) or by empty vector (C). (**II**) Evaluation of anthocyanin transcripts in *N. tabacum* (Xanthi) leaves following VIGS treatment: Wild type (A), treated with *pTRV2*:*VlmybA1-2* (B and D), treated with the empty vector (C, E and F). Expression of an endogenous actin gene (bp) was used as a positive control for RNA quality for all RNA samples.

**Figure 2.**
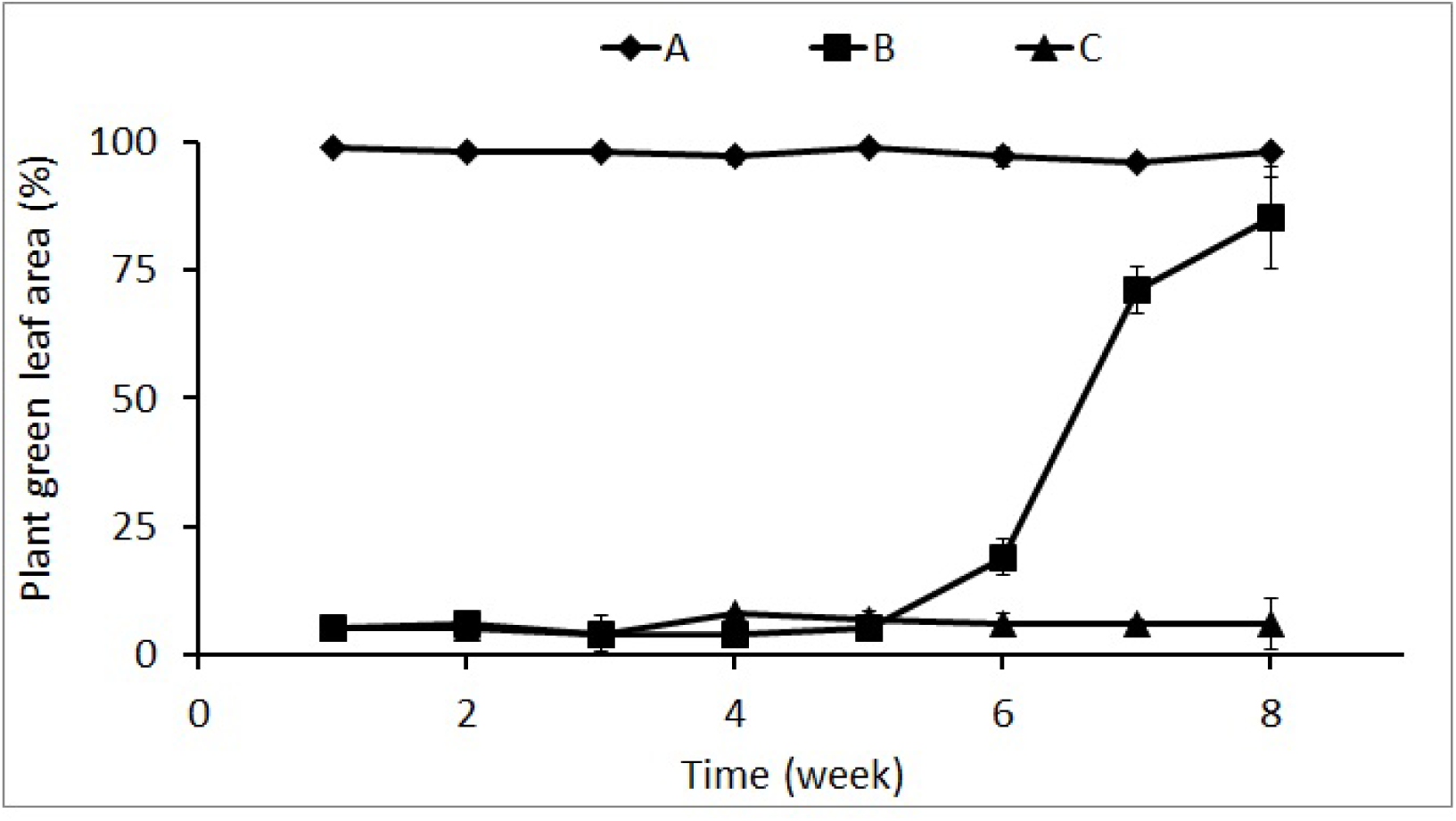
Effect of anthocyanin expression on the phenotypic expression of “green” leaves in wild type *tabacum* (Xanthi) plants (A), treated with pTRV2:*VlmybA1-2* (B), and treated with the empty vector (C).

**Figure 3.**
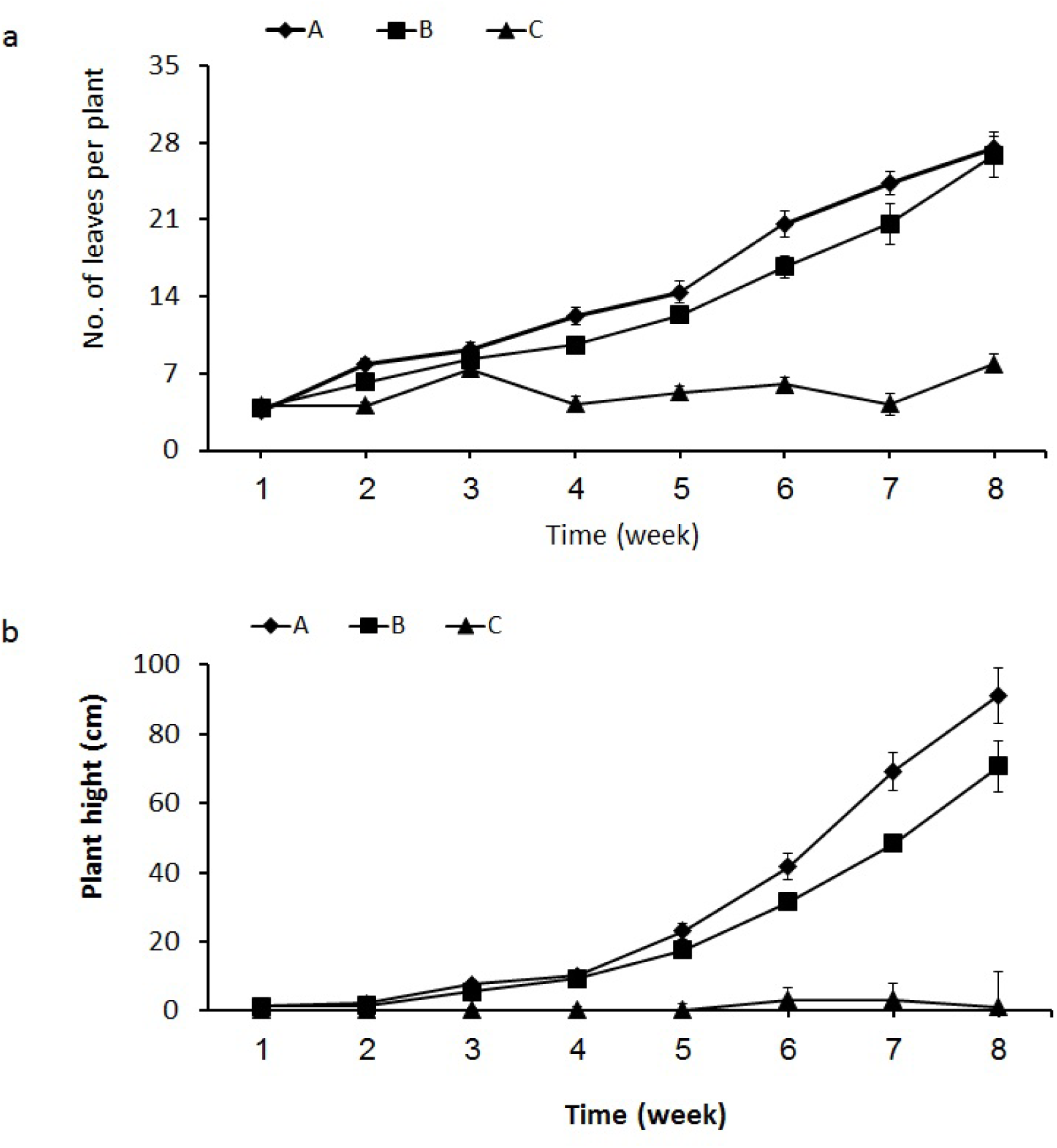
Effect of anthocyanin expression on leaves number (**a**) and height (**b**) of wild type *tabacum* (Xanthi) plants (A), treated with pTRV2:*VlmybA1-2* (B), and treated with the empty vector (C).

The transgenic “purple”-leaved plants developed in this study can promote PWM by overcoming the challenging task of weed/crop classification, using simple algorithms and highly accessible devices like a consumer-grade camera (3,7). The main contribution of this study is the ability to limit the anthocyanin gene’s expression to early growth stages, thereby avoiding a fitness cost. To make this technology applicable, we need another experimental system to apply or spray double-stranded (ds) RNA on the transgenic plants expressing the anthocyanin gene. Tenllado et al. (17) developed an in-vivo expression system to produce a large amount of virus-derived dsRNA in RNase III-deficient bacteria, with a view to providing practical control of a viral disease in plants. More recently, Wang et al. (18) reported that applying external siRNA or dsRNA that targets *Botrytis DCL1* and *DCL2* genes on the surface of fruit, vegetables and flowers significantly inhibits gray mold disease. Therefore, the discovery of RNA uptake by several organisms can be efficiently adapted for anthocyanin-derived dsRNA or sRNA for implementation in a field crop tagged with anthocyanin pigment. Adoption of the genetic modification and silencing concepts presented in this study may contribute to facilitating the use of PWM strategies and reducing herbicide usage, toward sustainable weed-control management.

## Methods

### Overexpression and silencing of *VlmybA1-2* in tobacco plants

For anthocyanin expression, we used the pBI121 vector carrying the full-length cDNA of the *VlmybA1-2* coding region under the control of the *CaMV 35S* promoter (9). Tobacco (*Nicotiana tabacum* cv. Xanthi) was transformed as described by Hamamouch et al. (19). Several kanamycin-resistant tobacco plants were selected for molecular and phenotypic analyses. For VIGS silencing of *VlmybA1-2*, a unique N-terminal fragment of about 146 bp was selected and amplified on the coding region of *VlmybA1-2* (Accession no. AB073012) on the pBI121 plasmid. The above fragment was amplified using the forward primer: 5’-CTGGTTCCCCTCCGAGCAGGGTTG-3’ overhanging the restriction site *XbaI* and reverse primer: 5’- GCAGTCCTCCCTGGAAGCCTACCCGC-3’ overhanging the BamHI restriction site. The gene construct was cloned into *pTRV2* at the respective sites and designated *TRV2*:*VlmybA1-2*. An empty *pTRV2* was used as the control treatment.

### VIGS assay and RT-PCR

The VIGS assay was performed on the F1 generation of transgenic tobacco (*N. tabacum* Xanthi) seedlings (two to three leaflets) overexpressing anthocyanin pigments. Agroinfiltration of tobacco leaflets was performed as described by Bachan and Dinesh-Kumar (20). A mixture of *Agrobacterium* cultures containing pTRV1 (RNA1) and p*TRV2*:*VlmybA1-2* at a 1:1 (v/v) ratio was injected into the lower surface of the tobacco leaflets. Total RNA was extracted from control and transformed tobacco plants using the TRI-REAGENT Kit (Molecular Research Center, Inc., Cincinnati, OH, USA). Systemic infection of pTRV1 and p*TRV2*:*VlmybA1-2* in leaves of *N. tabacum* plants was detected as described previously (14). To measure transcript levels of anthocyanin, semi-quantitative RT-PCR of *N. tabacum* wild type (WT; control), *pTRV2*:*VlmybA1-2* and empty *pTRV2* was performed (18 to 20 days after agroinfiltration). Total RNA (1 µg) was reverse-transcribed and the resulting cDNA was amplified by PCR using *VlmybA1-2*-specific primers 5’-CTGGTTCCCCTCCGAGCAGGGTTG-3’ (forward) and 5’-GCAGTCCTCCCTGGAAGCCTACCCGC-3’ (reverse).

To follow the F1 lines’ development, pot experiments were performed, aimed at quantifying their phenotypic expression and fitness. Wild type tobacco line was used as a reference for the transgenic lines expressing anthocyanin with respect to development and leaf hue. Five 2-L pots were used separately for each line: B (transgenic with silencing), C (silencing with empty vector) and A (WT). Plants were imaged once a week for 8 weeks using a conventional consumer-grade camera (Nikon D70 with Nikor 24 mm lens). The “green”-related parts of each plant were extracted as described by Lati et al. (21) and their relative area was visually estimated, followed by height and leaf number measurements. For each imaging time, percent of green leaf area (first experiment), height and leaf number were subjected to ANOVA (Tukey-Kramer Honestly Significant Difference test, *P* = 0.05). One-way ANOVA was conducted to determine the impact of the different lines on the estimated green leaf area, height and leaf number.

### Competing Interests statement

Prof. Radi Aly, Dr. Ran Lati, Vinay K. Bari, Mrs. Jackline Abu-Nassar and Prof. Hanan Eizenberg declare no potential conflict of interest.

All authors, have agreed to its submission, are responsible for the contents and have agreed that the corresponding author may sign on behalf of the other authors.

### Author Contributions

A.R. conceived, planned and supervised the work. L.A., V.K.B. designed and performed the experiments and analyzed the data. A.J. and E.H. contributed in data production.

